# Experimental evolution supports signatures of sexual selection in genomic divergence

**DOI:** 10.1101/2020.09.07.285650

**Authors:** R. Axel W. Wiberg, Paris Veltsos, Rhonda R. Snook, Michael G. Ritchie

## Abstract

Comparative genomics has contributed to the growing evidence that sexual selection is an important component of evolutionary divergence and speciation. Divergence by sexual selection is implicated in faster rates of divergence of the X chromosome and of genes thought to underlie sexually selected traits, including genes that are sex-biased in expression. However, accurately inferring the relative importance of complex and interacting forms of natural selection, demography and neutral processes which occurred in the evolutionary past is challenging. Experimental evolution provides an opportunity to apply controlled treatments for multiple generations and examine the consequent genomic divergence. Here we altered sexual selection intensity, elevating sexual selection in polyandrous lines and eliminating it in monogamous lines, and examined patterns of divergence in the genome of *Drosophila pseudoobscura* after more than 160 generations of experimental evolution. Divergence is not uniform across the genome but concentrated in “islands”, many of which contain candidate genes implicated in mating behaviours and other sexually selected phenotypes. These are more often seen on the X chromosome, which shows divergence greater than neutral expectations. There are characteristic signatures of selection seen in these regions, with lower diversity and greater *F*_ST_ on the X chromosome than the autosomes, and differences in diversity on the autosomes between selection regimes. Reduced Tajima’s D implies that selective sweeps have occurred within some of the divergent regions, despite considerable recombination. These changes are associated with both differential gene expression between the lines and sex-biased gene expression within the lines. Our results are very similar to those thought to implicate sexual selection in divergence in natural populations, and hence provide experimental support for the likely role of sexual selection in driving such types of genetic divergence, but also illustrate how variable outcomes can be for different genomic regions.

**Impact Summary:** How does sexual selection contribute to the divergence of genomes? It is often thought that sexual selection is a potent force in evolutionary divergence, but finding ‘signatures’ of sexual selection in the genome is not straight-forward, and has been quite controversial recently. Here we used experimental evolution to allow replicate populations of fruit fly to evolve under relaxed or strengthened sexual selection for over 160 generations, then sequenced their genomes to see how they had diverged. The features we find are very similar to those reported in populations of natural species thought to be under strong sexual selection. We found that genomic divergence was concentrated in small patches of the genome rather than widespread. These are more often seen on the X chromosome, which overall shows especially elevated divergence. There are also characteristic signatures of selection seen in these regions, with lower genetic diversity suggesting that selection was strong in these regions. The changes are associated with both differential gene expression between the lines and sex-biased gene expression within the lines. Many of the patches of divergence also contain candidate genes implicated in mating behaviours and other sexually selected phenotypes. Our results provide experimental support for the likely role of sexual selection in driving such types of genetic divergence.

## Introduction

The role of sexual selection in influencing evolutionary divergence and speciation is unclear (Panhuis *et al*., 2001; Ritchie, 2007; Maan & Seehausen, 2011; Servedio & Boughman, 2017). Associations between species diversity and proxies of sexual selection such as sexual dimorphism or mating system variation often imply that sexual selection can accelerate divergence, especially when acting alongside natural selection (Arnqvist *et al.*, 2000; Gage *et al*., 2002; Ellis & Oakley, 2016). However, different indicators of sexual selection give contrasting results in such comparative studies, and a consensus is not clear (Kraaijeveld *et al*., 2011; Janicke *et al*., 2018). One potentially compelling source of evidence that sexual selection is involved in divergence is coming from the increasing number of comparative genomic studies available across a range of organisms. Many descriptions of genomes, including those of species thought to have undergone strong sexual selection such as the Hawaiian *Drosophila* or African cichlids, have found that genes associated with mating behaviour or sensory perception potentially involved in sexual communication are often outliers in measures of divergence (e.g. Mattersdorfer *et al*., 2012; Kang *et al*., 2016). It has also been known for some time that genes which diverge particularly rapidly and show stronger signatures of positive divergent selection are often sex-biased in expression (Pröschel *et al*., 2006; Ellegren & Parsch, 2007; Zhang *et al*., 2007). Sex-biased gene expression itself, especially male-biased expression, evolves rapidly and this is associated with indicators of sexual selection such as increased sexual dimorphism in birds (Harrison *et al*., 2015; Wright *et al*., 2019). However, genes with sex-biased gene expression might experience more drift than unbiased genes, either due to reduced pleiotropy (Gershoni & Pietrokovski, 2014; Allen *et al*., 2018) or because they experience only half the selection pressure of genes with unbiased expression (Dapper & Wade, 2020). Additionally, divergence of sex chromosomes between species is usually much greater than autosomes, sometimes dramatically so (Counterman *et al.*, 2004; Ellegren *et al*., 2012).

However, such patterns of divergence are not necessarily driven by elevated sexual selection on these genes or genomic regions. Sex-biased gene expression is thought to evolve due to sexually antagonistic selection on gene expression, which is an important factor in sexual selection but can arise due to other types of conflict. Changes in sex-bias in gene expression are also complicated by additional factors including dosage compensation, turnover of sex-biased expression and resolution of conflict via sex-linkage or sex-limited expression (Mank *et al*., 2010a; Wright *et al*., 2019). The increased divergence of sex chromosomes is also potentially influenced by many factors, including a greater role of genetic drift due to a smaller effective population size on X chromosomes compared to autosomes, dominance effects, and other consequences of sex-linkage such as dosage compensation (Vicoso & Charlesworth, 2006; Ellegren, 2009; Mank *et al*., 2010b). Hemizygosity results in a lower effective population size (N_e_) on the X (N_eX_) than on autosomes (N_eA_). Under random mating the ratio of N_e_ is expected to be 3:4 and this should reduce neutral diversity and increase between-species divergence by the same proportion (Vicoso & Charlesworth, 2006). Hemizygosity should also result in an increased efficacy of selection for partially recessive beneficial mutations on the X-chromosome, relative to autosomes, and against recessive deleterious mutations on the X, relative to autosomes. Finally, because of the female-biased inheritance patterns of X-linked loci (males transmit them only to daughters while females transmit them to both daughters and sons), sex-limited selection as well as sexual selection will influence their divergence (Mank *et al*., 2010a; Corl & Ellegren, 2012; Wright *et al*., 2015).

It is very difficult to infer the historical role of different evolutionary processes from patterns of contemporary divergence between populations and species, because they can result in similar genomic signals (Butlin *et al*., 2012). One way of directly addressing the role of sexual selection or mating system variation in genomic divergence is to examine the genomic consequences of experimental evolution under manipulated sexual selection regimes in the laboratory. A great advantage of this approach is that there are potentially fewer confounding variables involved than when making comparisons across species or natural populations. However, a disadvantage is that the time scale over which divergence can be studied is typically much shorter than evolutionary time-scales in nature. Studies of experimental evolution and speciation are in their infancy, and general conclusions are, as yet, difficult to draw (White *et al*., 2020). Enforcing monogamy in otherwise polyandrous species will lead to both changes in the intensity of sexual selection and the balance of sexual conflict, as it effectively eliminates sexual selection and sexually antagonistic selection. A classic example of such manipulation is where *D. melanogaster* were kept under enforced monogamy for about 50 generations (Holland & Rice, 1999). Females from the monogamy treatment had reduced longevity compared to ancestral females, when exposed to ancestral males. This was expected because the reduction of conflict should favour less harmful males and females that are less resistant to male harm. Other experimental evolution studies under altered mating systems have been performed in dung flies (Hosken *et al*., 2001; Hosken & Ward, 2001; Martin & Hosken, 2003), different species of fruit flies (*D. melanogaster;* (Gerrard *et al*., 2013; Hollis *et al*., 2014; Innocenti *et al*., 2014; Perry *et al*., 2016); *D. pseudoobscura;* (Crudgington *et al*., 2005); *D. serrata;* (Chenoweth *et al*., 2015), seed beetles (McNamara *et al*., 2020) and hermaphroditic flatworms (Janicke *et al*., 2016). Though aspects of the treatments differ amongst such experiments, some common patterns have emerged. Gene expression changes are seen, especially of genes that are initially sex-biased, though the details can vary between studies (Hollis *et al*., 2014; Veltsos *et al*., 2017). Moreover, gene expression changes can be more pronounced for genes expressed in reproductive tissues (Innocenti *et al*., 2014), and genes involved in the post-mating physiological manipulation of female egg-laying and re-mating rates (Perry *et al.*, 2016).

A feature emerging from genomic comparisons between diverging species is that details of genomic architecture complicate the assessment of patterns of divergence across chromosomes. Whole chromosomal regions can show correlated responses due to reduced recombination and hitchhiking effects, especially in species with segregating inversions. Early studies of species differences interpreted “islands” of divergence in the genome as resulting from divergent selection on genes within these regions with gene flow homogenising the genetic background (Turner *et al*., 2005; Nosil *et al*., 2009). More recently it has been appreciated that chromosomal inversions and other regions of low recombination or diversity can accentuate such clustered divergence (Noor & Bennett, 2009; Cruickshank & Hahn, 2014; Wolf & Ellegren, 2016; Ravinet *et al*., 2017). “Barrier loci”, genomic regions under divergent selection that restrict gene flow (Butlin & Smadja, 2018), may occur within such clusters but the lack of recombination makes them difficult to localise precisely. In experimental evolution the amount of recombination will be determined by both genomic architecture and the number of generations completed during the study, which is often modest in studies of eukaryotes. Also, in experimental evolution the lines can be kept effectively allopatric, so homogenising gene flow in regions not experiencing selection should be absent. The genomic divergence which occurs during experimental evolution is usually extensive, with widespread differences dispersed throughout the genome (Kawecki *et al*., 2012; Tobler *et al*., 2014; Michalak *et al*., 2019).

Here we directly test the influence of sexual selection on genomic divergence. We examine replicated experimentally evolved lines of *D. pseudoobscura* in which sexual selection has been manipulated for over 160 generations. One set of 4 replicate lines were raised under enforced monogamy (M lines), which should eliminate both sexual selection and conflict. Another 4 replicates were reared under elevated polyandry (E lines), with 6 males per female. Polyandry mediates the strength of both intra- and intersexual selection and sexual conflict (Pizzari & Wedell, 2013) and elevated polyandry will increase both pre- and post-copulatory sexual selection via female choice and sperm competition beyond levels experienced in most natural populations (Snook, 2014). Previous studies of these lines have found divergence in some, but not all, of the types of traits predicted to diverge under sexual selection. Sperm morphology and heteromorphism, and testis mass did not diverge, but E males had larger accessory glands and a greater mating capacity (Crudgington *et al*., 2009), were more competitive in mating encounters (Debelle *et al*., 2016), and produced more attractive courtship song than M males (Debelle *et al*., 2017). Coevolutionary changes have occurred in female song preferences (Debelle *et al*., 2014). Sexually dimorphic cuticular hydrocarbons have also diverged between the lines (Hunt *et al*., 2012).

Patterns of gene expression have also changed between the lines. E females show an increase in expression of genes normally enriched in ovaries (Immonen *et al*., 2014). Sex-biased genes responded more strongly to the sexual selection treatment, but the direction of gene expression changes differed between sexes, tissues, and according to courtship experience (Veltsos *et al*., 2017). In most cases, the transcriptome was “feminised” under polyandry (i.e. female-biased genes were up-regulated or male-biased genes down-regulated in E lines), in a striking contrast to a similar study with *D. melanogaster* (Hollis *et al*., 2014). Males changed in patterns of gene expression in the testes and accessory glands, and changes in gene expression in females following mating also diverged, especially in the female reproductive tract (Veltsos et al. *in prep.).*

Here we examine genomic divergence between these lines using a pool-sequence approach (Schlötterer et al., 2014) after more than 160 generations of experimental evolution. The relatively long time-scale of this study should reduce linkage effects on allele frequency changes. We adopt a statistical approach that identifies alleles that have changed in frequency consistently across the replicates, to help reduce the potentially confounding effects of drift or replicate-specific selection (Wiberg et al. 2017). We find that divergent SNPs are not distributed randomly across the genome, but occur in distinct, obvious clusters. We examine what genes are involved and find several with mutant phenotypes related to mating and courtship behaviours. We found that the X chromosome has accumulated more divergence than the autosomes and explore if divergence is associated with recombination rate or changes in gene expression between the experimental lines.

## Methods

### Experimental Evolution

A full description of the experimental evolution procedure is available elsewhere (Crudgington *et al*., 2005). Briefly, a population of *D. pseudoobscura* was established from 50 wild caught females, bred in the laboratory for four years then four independent monogamy (M) and elevated polyandry (E) lines were established. M females were housed with a single male and E females with 6 males, with females typically mating with two or three males. The effective population size was maintained around 120 (Snook *et al*., 2009) for both treatments to try to minimise confounding effects of drift and treatment. At each generation, offspring were collected and pooled together for each replicate line, and a random sample used to constitute the next generation in the appropriate sex ratio, thus reflecting the differential offspring production across families (Crudgington *et al*., 2005; Crudgington *et al*., 2009). Enforced monogamy is expected to eliminate sexual selection and sexual conflict while elevated polyandry increases both pre- and postmating sexual selection and sexual conflict beyond levels encountered in most natural populations and in the ancestral population (Crudgington *et al*., 2005; Bacigalupe *et al*., 2007; Crudgington *et al*., 2009).

### Sequencing and Mapping

Sequencing was carried out after ca. 160 generations of selection (specifically, 164 for replicate 1, 163 for replicate 2, 162 for replicate 3, and generation 160 for replicate 4). Two pools of 40 females (one E and one M) were taken from each replicate line and genomic DNA extracted using a standard Phenol-Chloroform extraction protocol. Each pool was sequenced across two lanes on a Illumina HiSeq platform at the Center for Genomic Research (CGR) at the University of Liverpool. Details of coverage are provided in the Supplementary Material. Reads from each sequenced pool were mapped to the *D. pseudoobscura* reference genome (FlyBase v3.1 February 2013) using BWA mem (v. 0.7.7; Li, 2013). Alignments were filtered to remove duplicate reads, reads with a mapping quality < 30, and any reads which were not properly paired, using samtools (v 1.3; Li *et al*., 2009 following Schlotterer *et al*., 2014). Reads were locally re-aligned around indels using GATK (v3.7.0; McKenna *et al*., 2010; DePristo *et al*., 2011). The.bam files for each line were then merged using bamtools (Barnett *et al*., 2011) and the genome-wide coverage calculated from these merged files with bedtools (v. 2.26; Quinlan & Hall, 2010). SNPs were called using a heuristic SNP calling algorithm (PoolSNP; Kapun *et al*., 2020). Sites were considered only if the total coverage at the site was > 17 and < the 95^th^ percentile for each contig or chromosome. An allele was only called if the count for that allele across all pools was > 16 and the allele frequency across all pools was > 0.001. Nearly 2 million SNPs were called and used in downstream analyses (see Supplementary Material).

### Genomic Analyses

#### Identifying Consistent Allele Frequency Differences

Many evolve and resequence studies of *Drosophila* find that a multitude of SNPs have diverged, perhaps tens of thousands (Michalak *et al*., 2019). The number is inflated upwards at least in part due to segregating inversions and other areas of low recombination, and hitchhiking (Barghi & Schlotterer, 2019). In order to focus on the loci most likely to have diverged due to the treatment, we only considered as candidate SNPs those which diverged consistently across all 4 replicate pairs of lines. We identified these using quasibinomial Generalised Linear Models, which are less prone than other statistical approaches to be influenced by strong divergence in only some replicates (Wiberg *et al*., 2017). The model structure applied was;

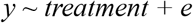

where *y* is the allele frequency of the major allele (identified as the major allele across all pools) within each sample, *treatment* is the experimental evolution treatment regime of each sample, and *e* is a quasibinomially distributed error term. If any count within a population was 0, +1 was added to all counts. P-values were converted to q-values using the “qvalues” R package (v. 2.16.0; Storey & Tibshirani, 2003). A threshold of 0.05 was chosen to control the false discovery rate (FDR), thus we define “top SNPs” as those which change consistently across all replicates with q-value < 0.05 and the remainder are referred to as “background” SNPs.

#### Genetic Diversity and Differentiation

We calculated genome-wide genetic diversity statistics (π and Tajima’s D) for windows of 50kb (with a 10kb overlap) using available python scripts (Kapun *et al*., 2020). Similarly, we computed pairwise *F*_ST_ estimates between E and M line pairs for each SNP using the R package “poolfstat” (v. 0.0.1; Hivert *et al*., 2018), averaged in windows of 50kb (with a 10kb overlap between windows). Comparisons of parameters between selection regimes and genomic regions were tested using non-parametric Wilcoxon tests. Additionally, we estimated neutral expectations for *F*_ST_ expected from drift and differences in effective population sizes on X chromosomes (*F*_X_) as in (Machado *et al*., 2016) using the equations of (Ramachandran *et al*., 2004) (equation 8 therein), *F*_X_ is given by:

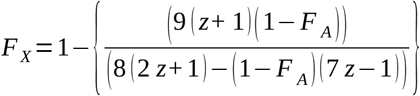

where, *z* is the ratio of the number of breeding males to females and *F*_A_ is the observed *F*_ST_ on autosomes. We assumed *z* to be either 1 or 6 to represent extreme possibilities based on the mating system manipulation. For each E-M pairwise comparison, we calculated mean *F*_ST_ across each chromosome type and converted to *F*_X_. We used a bootstrapping approach to obtain a random distribution of *F*_X_ for each replicate. For each of 1,000 bootstrap iterations we sampled, with replacement, a number of windows equal to the total number across all autosomes from the set of all windows, then we calculated mean *F*_ST_ across all sampled windows and converted to *F*_X_ using the equation above. Additionally, we computed a value of *F*_ST_ and Tajima’s D for each annotated *D. pseudoobscura* gene by taking the mean value across all 50kb windows that spanned a gene.

#### Linkage Disequilibrium (LDx)

Although haplotype information is not available from pool-seq data, short range linkage information is available from paired reads. We used LDx (Feder *et al*., 2012) to first compute the r^2^ of SNPs located on the same read pairs. We only used SNPs with a minor allele frequency > 0.1, a minimum coverage of 10, a maximum read coverage of 400, and a phred score > 20. Note that the empirical median insert size varied between 332-346 across samples. We binned pairs of SNPs into distance classes and then computed mean r^2^ per distance class. We only used distance classes with a minimum of 5 SNPs. We estimated the decay of r^2^ as a function of distance by fitting a linear model of r^2^ as a function of the log of the distance between the SNPs. Thus, the slope measures the decay rate of linkage due to recombination (Feder *et al*., 2012), giving an indication of the distance over which LD is present. In regions of low recombination one would expect high overall values of r^2^ but a weakly negative slope as LD is maintained over relatively longer regions of the genome. Comparing the slope parameter across different genomic regions gives an indication of differences in the recombination rate (or extent of selective sweeps). This was performed for each chromosome, as well as for different regions on the 3^rd^ chromosome (see below).

#### Functional Genomics

To examine the function of genes near candidate SNPs we conducted enrichment analyses. We used the *D. pseudoobscura* annotation and a dataset of regulatory long noncoding RNAs (lncRNAs; Nyberg & Machado, 2016). We identified genes or lncRNAs within a distance of 10kb up- or downstream of top SNPs with bedtools (Quinlan & Hall, 2010) intersect (keeping any potential ties). Enhancer regions, transcription factor binding sites, and other regulatory regions can occur up to 1 Mb up- or downstream from a target gene in other species (e.g. Maston *et al*., 2006; Chan *et al*., 2010; Werner *et al*., 2010; Pennacchio *et al*., 2013) but typically lie within 2kb of a gene region in *D. melanogaster* (Arnosti, 2003), 10kb thus represents a compromise. We submitted the implicated genes to ModPhEA (Weng & Liao, 2017) for phenotypic enrichment analysis. We combined the phenotypic classes “courtship behavior defective” (FBcv:0000399) and “mating rhythm defective” (FBcv:0000401) into one phenotype group and also tested the phenotypic class “stress response defective” (FBcv:0000408) for enrichment. We chose these classes *a priori* because they were most likely to be involved in phenotypic differences between the treatments related to mating or courtship behaviour and responses.

We also took advantage of gene expression data from the same experimental evolution lines. Expression data is available from heads and abdomens of virgin and courted flies (Veltsos *et al*., 2017) and testes, accessory glands, ovaries and female reproductive tracts from virgin flies, and ovaries and female reproductive tracts from mated females (Veltsos et al., *in prep.).* Using these data we compiled a list of genes with differential expression between E and M lines. For simplicity we considered a gene to be differentially expressed between E and M lines if it shows significant differences in E/M contrasts in any of the following data: combined virgin and courted head or abdomens of each sex (4 sets), virgin individual reproductive tissues (4 sets), mated individual female reproductive tissues (2 sets). Briefly, the analysis was conducted in edgeR v3.18.1 (Robinson *et al*., 2010) running in R v.3.4.0 (R Development Core Team, 2007). We used TMM normalization in edgeR and measured dispersion using a negative binomial model from the genes within each contrast. We employed a statistical definition for differential expression (FDR < 0.05; (Benjamini & Hochberg, 1995) and did not require a minimum logFC threshold to consider a gene differentially expressed as the effect of allometry should be minimal for samples from specific organs (Montgomery & Mank, 2016), and the results are cross-checked with top SNPs, making the analysis conservative. The associated scripts and final #number# gene set are available in OSF1, OSF2, File S#.

We used this list to ask if top SNPs co-localised with genes that are differentially expressed between the lines and if these also show different levels of diversity (Tajima’s D) or differentiation (*F*_ST_) between E and M lines. We used a resampling approach, sampling genes (without replacement) from the *D. pseudoobscura* annotation, to determine the amount of overlap with the DE genes that is expected by chance. For each sample, we picked a set of 428 genes from the annotation, which is the same size as the set of genes near top SNPs (see Results). We then calculated the proportion of these genes that also occur in the DE gene sets and repeated this procedure 1,000 times to build a distribution of expected overlap between re-sampled gene-sets and the DE gene sets. If the empirical set of genes near top SNPs had a proportional overlap >= the 95^th^ percentile of the re-sampled distribution it was deemed a “significant” overlap.

Using the values of Tajima’s D and *F*_ST_ computed for each gene (see above) we also asked whether there was any evidence of different levels of diversity or divergence between DE genes in any set (N = 3,173) and non-DE genes (N = 13,583). For Tajima’s D we contrast DE and non-DE genes separately for each chromosome type (autosomes, X-chromosome left arm, X-chromosome right arm), and each experimental evolution treatment (E and M; 6 contrasts in total), using Wilcoxon rank sum tests. For *F*_ST_ we contrast DE genes and non-DE genes separately for each chromosome type (3 contrasts), testing for differences with Wilcoxon rank sum tests. In both cases, the mean value for non-DE genes was used as a single value against which to compare DE genes, which reduces the effect of the enormous sample size for the non-DE genes on the significance of the test.

Finally, we also asked whether the changes in sex-biased expression (data from Veltsos *et al*., 2017) between E and M treatments (ΔSB_EM_) was related to diversity (Tajima’s D) within either E or M lines. Sex-bias in expression was assessed for two tissues, head and abdomen, in both courted or virgin data combined. Within each tissue, sex-bias was computed as the log_2_(fold change) in expression between males and females in E and M lines separately, after which ΔSB_EM_ is calculated as log_2_(FC)_E_ – log_2_(FC)_M_. Thus, positive values of ΔSB_EM_ correspond to greater male-bias in expression in the E lines, while negative values correspond to greater male-bias in the M lines. ΔSB_EM_ was then related to values of Tajima’s D in either E (TajD_E_) or M (TajD_M_) lines. For each tissue (head and abdomen) we performed an ANCOVA with chromosome (autosome, X-chromosome right arm, and X-chromosome left arm) as a co-factor, as well as mean Tajima’s D across E lines and mean Tajima’s D across M lines as co-variates. We also included the interactions between Tajima’s D and chromosome. The full model is:

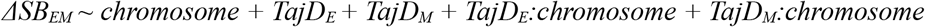

We further extracted the 30bp up- and down-stream of each SNP from the reference genome using gffread from the Cufflinks package (v2.2.1; (Trapnell *et al*., 2010) and tested for an enrichment of TF binding site motifs around top SNPs with the AME routine from the MEME package (v. 4.10.2; (McLeay & Bailey, 2010). GO term enrichment analysis was performed with GOwinda (v. 1.12; (Kofler & Schlotterer, 2012). We considered SNPs to be associated with genes if they occurred within 10kb up or downstream of an annotated gene. An empirical p-value distribution was produced from 1 million simulated SNP sets.

All statistical analyses were made with R (v. 3.6.3; R Development Core Team 2020) except where otherwise stated. Figures were drawn using the “ggplot2” package (v. 2.2.1; (Wickham, 2009) and associated packages (table S1).

## Results

### Consistent Allele Frequency Differences

In total, 480 SNPs show significant consistent allele frequency differences due to the experimental evolution treatment (hereafter the “top SNPs”). These occur on all of the main chromosomes but many show striking co-occurrence into a few clusters of highly differentiated SNPs (figure 1A). The distribution of the top SNPs across the genome is not random, with a significant excess on the 3 ^rd^ chromosome and both arms of the X chromosome (table S3). In particular, a large cluster of differentiated SNPs are observed at the end of the right arm of chromosome 3 (figure 1A). Other large clusters occur on both arms of the X chromosome (figure 1A). If all top SNPs within 50kb of others are grouped into clusters, this produces 70 distinct clusters throughout the genome (figure 1A). The majority of SNPs (72.9%) occur in only 6 clusters with > 10 SNPs.

**Figure 1.**
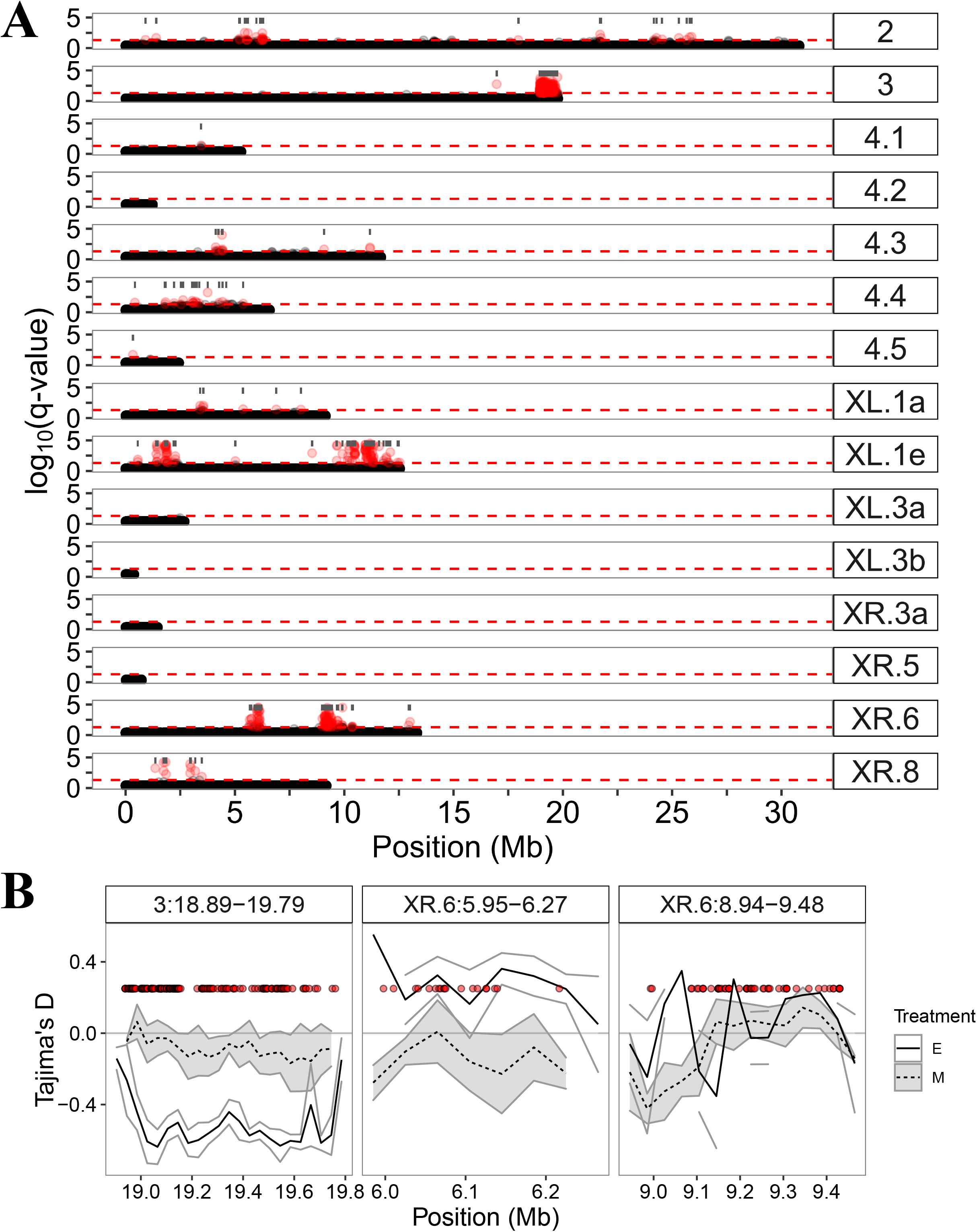
**A)** Manhattan plot of log10(q-values) for each SNP from a quasibinomial GLM with treatment as a predictor. Red points denote SNPs with a q-value < 0.05 and the horizontal red dashed line indicates the q < 0.05 cutoff. Grey bars give the locations and span of the 70 divergent regions (see text). **B)** Mean (± SE) Tajima’s D across replicates for the three most divergent regions (see text), red points denote SNPs with a q-value < 0.05, all have been plotted at the same value on the y-axis for convenience.

Such clustered divergence is often seen in comparisons between natural species (Ravinet *et al*., 2017) but rarely in experimental evolution (e.g. Kauranen *et al*., 2019). We considered 10 random permutations of the treatment labels among SNP sets and observed far fewer SNPs with q-values < 0.05 than in the original dataset. We are therefore confident that our approach reliably identifies SNPs with consistently different allele frequencies between the treatments. We also tested if the divergence was more clustered than random samples between the lines using a permutation test (for full details of the randomisation tests see the Supplementary Material). We also examined if variation in coverage might be associated with calling clustered divergence. We compared coverage within these clusters to 100 random genomic regions with a similar distribution in size shows that, although there is a minor difference in coverage between peaks with top SNPs, the variation in coverage across samples is far greater, we therefore conclude that difference in coverage around top SNPs and the rest of the genome cannot explain the patterns (figure S1).

The clusters do not correspond to known inversions in *D. pseudoobscura.* In particular, the large cluster on chromosome 3 containing many (N = 199, 41.5%) top SNPs does not correspond to the most common inversions that have shaped the evolution of this chromosome in the wild (Wallace et al., 2011; Wallace et al., 2013). Allele frequencies in E and M lines for the top 100 SNPs are shown in figure S2. More than half of these (57%) are fixed differences in all replicates. Across all the top SNPs, 12% are fixed differences between the E and M lines in all replicates, with all of these occurring on the X chromosomes

### Genetic diversity

We identified a set of candidate SNPs which varies consistently in allele frequency in response to experimental treatment. Such patterns are strongly suggestive of the action of selection. We therefore also assessed the levels of genetic diversity throughout the genome and in regions surrounding these candidates. On a broad scale, Tajima’s D does not vary much across chromosomes (figure S3). Strikingly, Tajima’s D is substantially lower on chromosome 3, though the interaction effect of chromosome and treatment is not statistically significant (F_4,30_ = 0.59, p = 0.68). Strongly localised selective sweeps, should locally reduce Tajima’s D. Within E lines, Tajima’s D is actually on average slightly higher within the clusters containing top SNPs (−0.03) than outside these clusters (−0.05; Wilcoxon signed rank test: V = 17623, p-value = 0.04). Within M lines there is no statistically significant difference between clusters (−0.07) and outside clusters (−0.06; V = 13390, p-value = 0.3). However, patterns of Tajima’s D are very variable. The most differentiated region on chromosome 3 shows reduced Tajimas’s D within the E treatment compared to the M treatment (figure 1B), as would be expected following selective sweeps. Similar patterns are seen for some peaks on the X chromosome (figure S4). In a few cases, there are reductions of Tajima’s D associated with regions containing top SNPs within M lines compared to E lines (figure 1B and figure S4). However, many of these regions are quite small and consequently estimates of Tajima’s D may be unreliable (figure S3).

Nucleotide diversity across the chromosomes was estimated as π (figure S5). Diversity is lower overall in E lines than in M lines (figure 2A). Diversity varies significantly across chromosomes in both E and M lines (figure 2A; F_4,30_ = 29.3, p < 0.001), but the interaction with treatment is not significant (F_4,30_ = 0.98, p = 0.44). Lowest diversity (in both treatments) is seen on the more differentiated chromosomes (X and 3; figure 2A). Median π is marginally non-significantly lower within the clusters of M (V = 12471, p = 0.05), but not E (V = 13843, p = 0.19), lines. The ratio of diversity between the sex chromosome and autosomes is lower in E lines than in M lines, though this is variable across replicates (figure 2B). Overall, it seems like there is greater evidence for selective sweeps in E lines, especially for the X.

**Figure 2.**
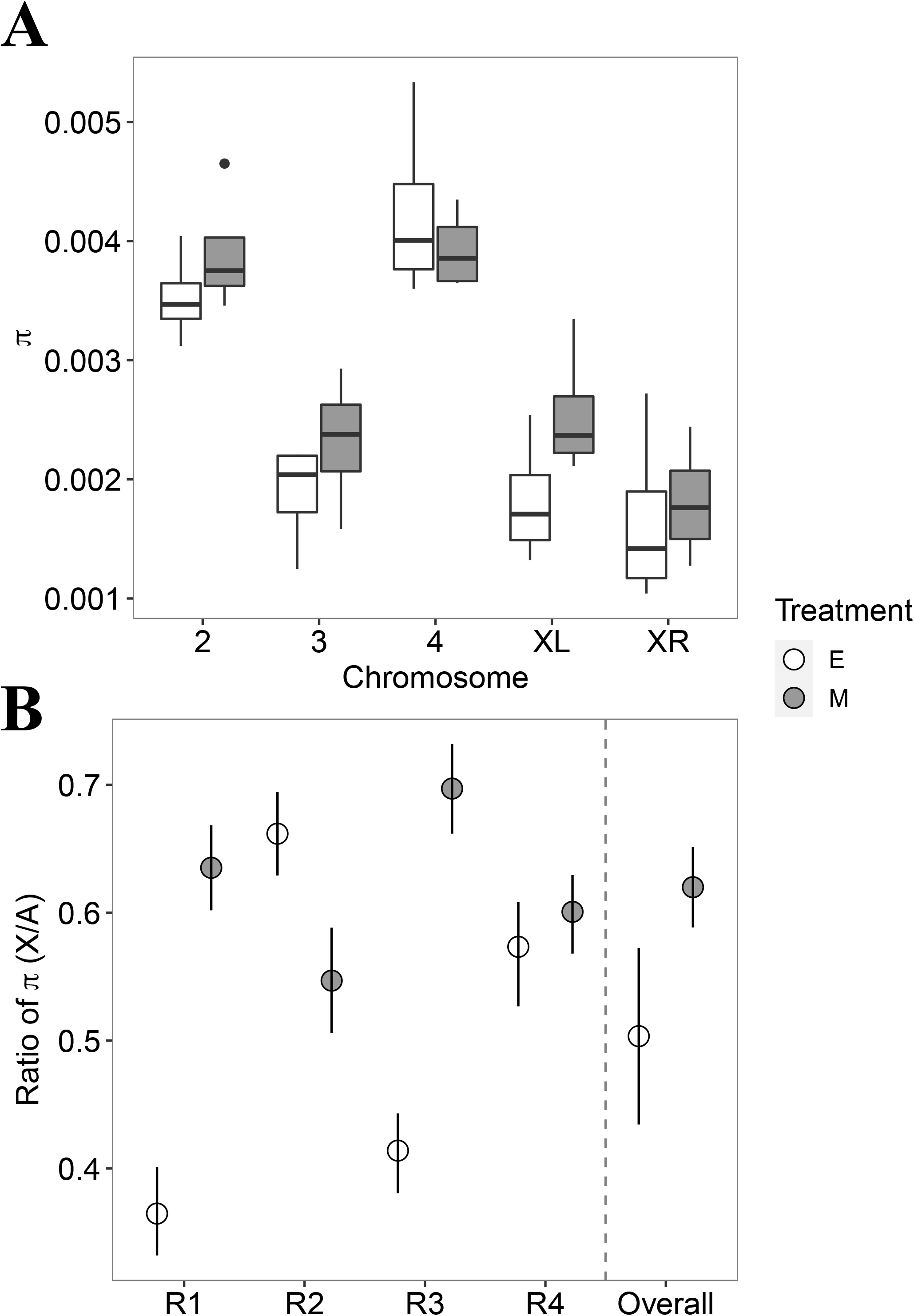
**A)** Levels of genetic diversity (π) on each chromosome in E and M lines. π is estimated in overlapping windows of 50kb, then averaged across the chromosomes. Boxplots show the distribution of π on each chromosome across replicate experimental evolution lines. **B)** The X chromosome to autosome ratio of π in the replicates of E and M lines and overall.

Comparisons of genomic divergence are often based on patterns of *F*_ST_. Although obviously not independent of changes in allele frequency, we also examined the patterns of *F*_ST_ seen between the E and M lines for comparison with published studies and to examine the X / autosome divergence in more detail. *F*_ST_ is generally higher on the X chromosome than on autosomes (figure 3B), even after accounting for the expected greater effects of drift on the X over the autosomes (see Methods for the equations; figure 3B). Hence the X:A ratio of *F*_ST_ is always > 1 (figure 3C). These results hold regardless of the value of *z* (see Methods for the equations). *F*_ST_ was higher within peak regions than outside peak regions (0.64 vs. 0.59; Wilcoxon signed rank test: V = 15309, p-value < 0.001, Figure 3D), as expected as allele frequencies differ most within the clusters. It should be noted that the above measures of differentiation and genetic diversity are often variable and precise estimates depend on the number of SNPs detected, the coverage, and number of replicate lines. Accordingly, we emphasise that while broad-scale patterns are likely to be robust, values for any one genomic region or gene should be taken with appropriate caution.

**Figure 3.**
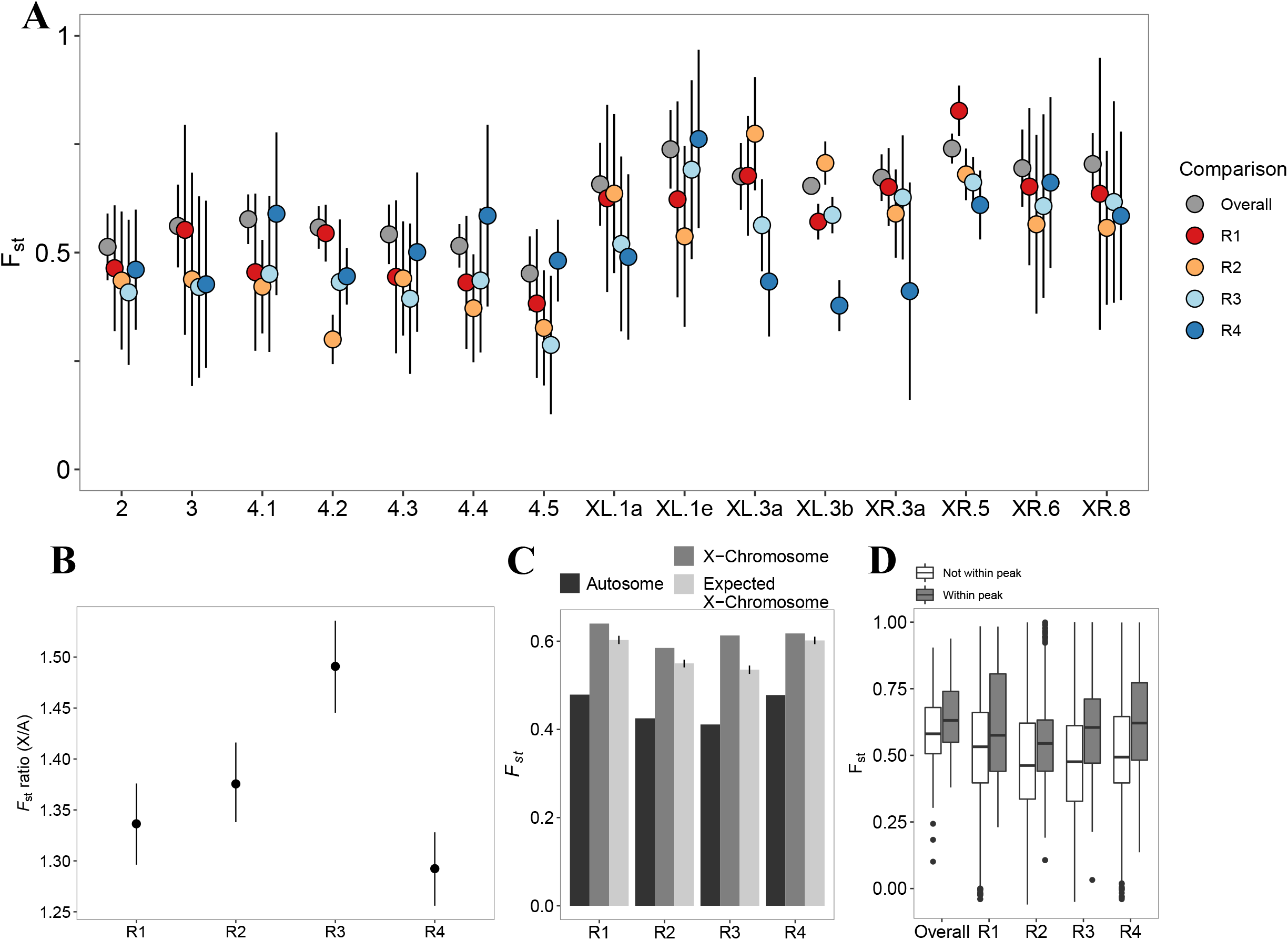
**A)** *F*_ST_ between E and M treatment lines on the main chromosome arms for each replicate. *F*_ST_ is calculated for each SNP then averaged within overlapping 50kb windows on each chromosomal segment. **B)** The X:autosome ratio of *F*_ST_ within each replicate line. The error bars are bootstrap 95% confidence intervals. **C)** Observed *F*_ST_ on the autosomes (black) and on the X chromosome (dark grey) as well as the expected *F*_ST_ on the X chromosomes assuming a value of *z* = 6 (light grey) (see Methods), error bars represent bootstrap 95% confidence intervals. **D)** The difference in FST between windows within “peaks” of top SNPs and windows outside of these peaks.

### Linkage Disequilibrium

Background selection or selective sweeps could lead to clustered genomic divergence, often with low diversity, especially in regions of low recombination such as telomeric regions. We examined patterns of linkage disequilibrium in the clusters and if this varied with treatment. Throughout the genome, the decay rate (*a* parameter) of LD is generally shallower (i.e. less negative) in the E treatment (figure 4A). This is seen for chromosome 3 as well as both arms of the X chromosome (figure 4A). A lower decay rate is indicative of more LD, due to less recombination and/or a potential for greater hitchhiking under positive selection. Contrary to predictions, we found a steeper rate of decay (less LD) within the differentiated region of chromosome 3 than outside it, especially in E lines (figure 4B and C). Although statistically significant (F_(2,13)_ = 4.6, p < 0.001), these differences are slight. The most striking pattern overall is greater overall LD on chromosome 3.

**Figure 4.**
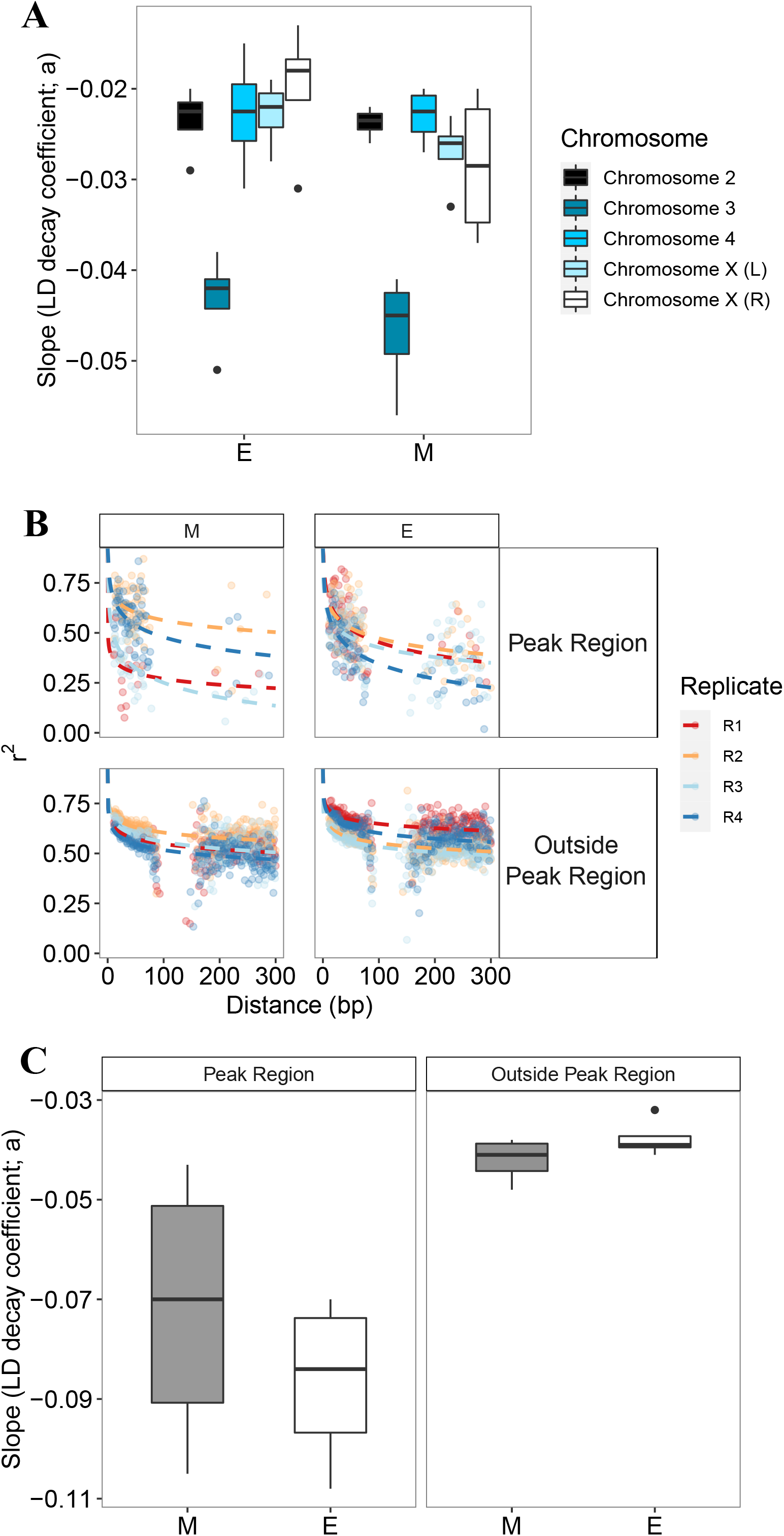
**A)** Slope coefficients from the model *r^2^ ~ a* + log(*bp*) where *bp* is the distance between pairs of SNPs and *r^2^* is the average measure of LD between SNPs. Distributions are shown for average values of each of the main chromosomes as well as X chromosomes across replicates in E and M lines. **B)** Decay in LD as a function of distance between SNPs with the chromosome 3 peak region (see figure 3) and outside the peak region for E and M lines. **C)** The distribution of slope parameters for SNPs within the chromosome 3 peak and outside the peak region.

### Gene functions and expression variation

Out of the 480 top SNPs, 201 (42%) lie within a gene model (i.e. either in an intron or within an exon; the remaining are intergenic. The top SNPs are not significantly enriched in any GO terms after correcting for multiple testing, even at a 10% FDR (table S4). Similarly, there is no enrichment of genes with annotations for mating behaviour or stress response phenotypic classes. However, several genes within 10kb of a top SNP are potentially interesting candidate genes for traits evolving under sexual selection based on described functions (table S4). For example, the genes *Odorant-binding protein 47a (Obp47), pickpocket 6 (ppk6),* and *Accessory gland protein 53C14c (Acp53C14c)* all occur within 10kb of a top SNP and are genes potentially underlying sexually selected behaviours or traits. Two of these genes *(ACP53C14c* and *Obp47a)* are within the region of highly differentiated SNPs on the 3^rd^ chromosomes, which also includes several additional accessory gland proteins *(Acp53Ea, Acp53C14b, Acp53C14a),* and other genes (table S4), all of which are thought to influence mating and courtship behaviours or phenotypes based on known functions of similar genes in *D. melanogaster*.

Previous studies have shown that there is divergence in gene expression patterns between E and M lines (Immonen *et al*., 2014; Veltsos *et al*., 2017; Veltsos et al., *in prep.).* We therefore asked if these expression differences were associated with the top SNPs. Genes within 10kb (N = 428) of the top SNPs show a significantly greater overlap with genes that are differentially expressed (DE) in ovaries and testes between E and M lines than expected by chance (figure S6 and table S3). This pattern also holds for genes within 1Mb (N = 7,045; figure S7). Also, there is evidence that *F*_ST_ between E and M lines is higher for genes that are DE between the lines, especially for X-linked genes (figure 5A; Wilcoxon rank sum tests, Autosomes - V = 1026000, p = 0.03; X-chromosome right arm - V = 89067, p = 0.005; X-chromosome left arm – V = 59623, p = 0.04). There is no evidence that Tajima’s D is different between DE and non-DE genes (Wilcoxon rank sum test; all p > 0.05; figure 5B). There is some evidence that the degree to which sex-biased expression of a gene changes between E and M lines is associated with Tajima’s D in M lines, but only on the X-chromosome and only within abdominal tissues (figure 5C). Specifically, as the change in sex-bias becomes more negative (i.e. more female-biased expression in M lines), Tajima’s D also becomes more negative (interaction of Tajima’s D in M lines and chromosome type: F_(11189,11191)_ = 4.4, p = 0.01).

**Figure 5.**
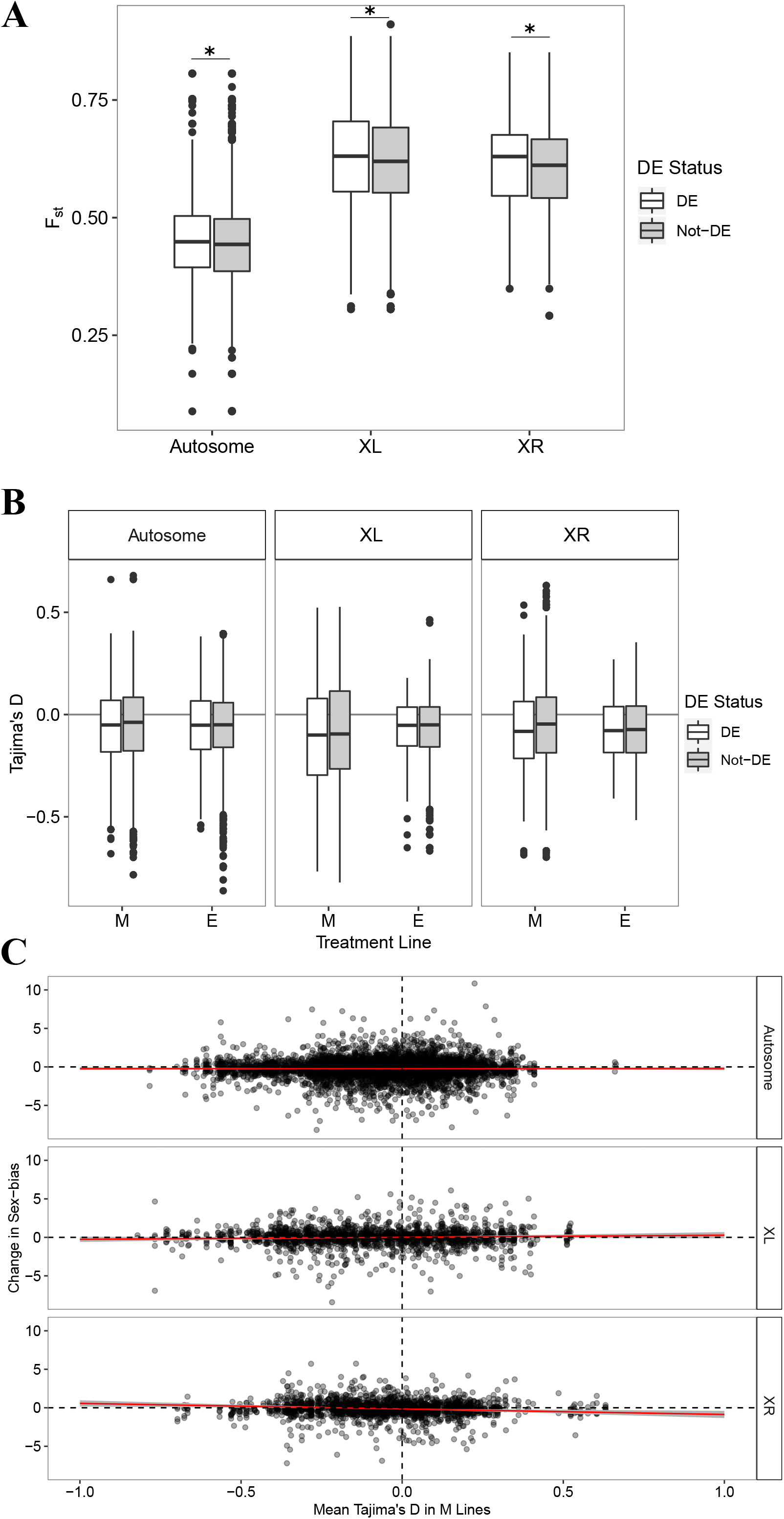
**A)** *F*_ST_ at DE vs. non-DE genes for different chromosome types. Asterisks indicate significant differences **B)** Tajima’s D at DE vs. non-DE genes for different chromosome types. **C)** relationship between change in sex-bias between E and M lines and Tajima’s D in M lines.

The regions immediately up- or down-stream of top SNPs are not enriched for TF binding motifs or lncRNAs, after correction for multiple testing, so there were no obvious differences between treatments in regions expected to influence gene expression variation.

## Discussion

There is much debate about the influence of sexual selection and sexually antagonistic selection on patterns of genomic variation (Mank, 2017; Sayadi *et al*., 2019) and how this may influence divergence between species (Wolf & Ellegren, 2016). Sex-biased gene expression, especially male-bias, evolves quickly and is related to phenotypic sexual dimorphism (Wright *et al*., 2019). Outliers in genome scans often implicate sexual selection as a diversifying force (Andres *et al*., 2008; Blankers *et al*., 2018). Sexual antagonism may be associated with genomic signatures of selective sweeps or balancing selection (Cheng & Kirkpatrick, 2016; Wright *et al*., 2019) and may be promoted by strong sexual selection (Connallon & Clark, 2012; 2013; Dutoit *et al*., 2018; Ruzicka *et al*., 2019). However, inferences of the sources of selection on natural variation in genomic divergence are usually indirect and ambiguous, because multiple forces act in concert to produce variation seen at the genomic level in nature. Here we used experimental evolution to alter sexual selection intensity, elevating sexual selection in polyandrous lines and eliminating it in monogamous lines, and examined patterns of divergence in the genome after more than 160 generations of experimental evolution.

Many of the results we found recapitulate patterns seen in natural populations and between species. Divergence is not uniform across the genome but clustered in “islands” of divergence, some of which contain candidate genes for an involvement in mating success. These clusters are more often seen on the X chromosome, which is a “hot spot” for divergence. There are signatures of selection within the islands of divergence, with marginally lower diversity (π) within clusters than the rest of the genome, but only in M lines. *F*_ST_ between E and M lines is greater within clusters, and is also greater on the X than autosomes, and differences in diversity are seen in the autosomes between selection regimes. Low Tajima’s D implies selective sweeps have occurred, but only within some of the divergent regions. These patterns of diversity and divergence are associated with changes in both differential gene expression between the lines and sex-biased genes. Overall, *F*_ST_ between the lines is high in all replicates, probably due to low overall effective population sizes, though effective population sizes are similar between E and M lines (Snook *et al*., 2009).

The concept of “islands” of divergence originated from comparisons of genomic divergence between species (Nosil *et al*., 2009; Ravinet *et al*., 2017). These are usually thought to have arisen due to the combination of strong selection on barrier loci and genetic hitchhiking within genomic regions, with background gene flow reducing divergence outside of the islands. Here we find distinct clustered divergence akin to the islands seen in natural systems. Our system is effectively allopatric, so there was no background gene flow counteracting divergence outside of these clusters, which therefore must have arisen due to strong localised divergent selection across all replicates. Although *D. pseudoobscura* has relatively well-characterised inversion polymorphisms (Sturtevant & Dobzhansky, 1936; Dobzhansky & Sturtevant, 1938; Wallace *et al*., 2011), the clusters we describe do not correspond to the most common inversions known for this species, which are often very large. Our short-read sequencing approach allowed some examination of LD and there was no suggestion of reduced recombination within the clusters. In fact, the large peak at the right end of chromosome 3 (figure 4) surprisingly seems to be within a region of high recombination (which is often suppressed at telomeric regions). Interestingly, recombination is higher within this peak than the chromosome-wide rate, but also differs between the treatments, being greater in the M lines. Perhaps selection against recombination was reduced in monogamous individuals because of epistatic interactions in the region which were important in sexual selection or sexual conflict. There was no obvious difference in LD in the other clusters but their smaller size and hence “noisier” estimates makes robust inferences from pool-seq data difficult. Indeed, the estimates of LD within the cluster on chromosome 3 also rely on relatively few SNPs at longer ranges compared to the rest of the chromosome, so inferences need to be taken with caution.

The lack of background gene flow or stronger linkage disequilibrium within the clusters suggests that they have arisen primarily through localised strong selection that is consistent across all replicates. In support of this, we see lower Tajima’s D in some of the larger clusters. However, these patterns are very variable with lower Tajima’s D in different clusters for the E and M lines. Thus, overall, there is no significant difference in Tajima’s D between E and M lines. Systematic differences in Ne between E and M lines might be expected to lead to consistent differences in Tajima’s D. One might predict lower Ne in M lines due to fewer mating individuals and, correspondingly, lower Tajima’s D in M lines, though the experimental design tried to minimise this and previous studies found no evidence of such a reduction in Ne (Snook *et al*., 2009.

The genes contained within the clusters are not enriched for genes of particular functional categories, however, they do include strong candidate genes for an involvement in mating system evolution. For example, the large region on chromosome 3 contains numerous accessory gland proteins. In *D. melanogaster* these are well known to influence male reproductive success, exert antagonistic effects on female fecundity and lifespan, and play a role in sperm competitive success (Chapman *et al*., 1995; Ram & Wolfner, 2007). Some of the evolutionary response in E lines is antagonistic, because M females have a lower fecundity when mated with E males. Moreover, when mated to E males, the reproductive schedule of M females is manipulated to the males benefit (Crudgington *et al*., 2010). Accessory gland proteins show accelerated coding sequence and expression evolution across species (Swanson & Vacquier, 2002; Begun & Lindfors, 2005). Other genes within the clusters are involved in sexual chemical communication, which is also often implicated in outlier analyses in genome comparisons between species (Smadja & Butlin, 2009). For example, mutants of members of the pickpocket family in *D. melanogaster* show aberrant male mating success because of their involvement in the detection of female pheromones (Thistle *et al*., 2012; Toda *et al*., 2012). E males, subject to both intra- and intersexual selection, have diverged in aspects of courtship behaviour, such as time until initiation of courtship, have a higher intensity courtship song and have a higher competitive mating success than M males (Debelle *et al*., 2016; Debelle *et al*., 2017).

If strong selection has driven this clustered genomic divergence, an interesting question is whether the responses to selection are stronger in the E or M lines. Imposing monogamy on a naturally polyandrous species probably leads to relaxed selection on many genes involved in intra-or intersexual competition. Therefore, the response is likely to involve changes in both the intensity and direction of selection on some loci. Thus, perhaps the variation in signals of selection we see in Tajima’s D and changes in LD are to be expected. Overall, we see stronger reductions in diversity in E lines, perhaps suggesting that directional selection was stronger when sexual selection was strengthened.

One pattern very commonly seen in studies of natural populations and species is more rapid divergence of the X chromosome (Vicoso & Charlesworth, 2006). We also see this here, the X having a higher prevalence of divergent clustered regions and consequently higher *F*_ST_ between the lines. Remarkably, all SNPs with fixed differences between the lines occurred on the X. Faster X evolution can occur for many reasons, including greater genetic drift due to its smaller effective population size, and beneficial recessive alleles on the X are more responsive to selection due to male hemizygosity (Meisel & Connallon, 2013). We calculated expected X/A divergence ratios under a range of plausible sex ratios and the observed X/A divergence exceeded all of them, suggesting the accelerated X divergence is not due to drift effects alone, selection or a combination of effects are likely involved. Genes under sexual selection are potentially more likely to be sex-linked, due to antagonistic, or sex-limited selection (Reinhold, 1998; Kirkpatrick & Hall, 2004). Sexually selected or antagonistic loci are perhaps also more likely to show dominance effects (Grieshop & Arnqvist, 2018).

Previously we found that gene expression differences have evolved between the lines, especially in sex-biased genes (Veltsos *et al*., 2017). Here we show that there is significant overlap between differentially expressed genes and the regions of genomic divergence of the lines found here. Thus, the expression divergence is associated with the broad patterns of genomic divergence. Also, *F*_ST_ is greater for the differentially expressed genes, once again recapitulating patterns from natural systems (sex-biased genes here are not more likely to be sex-linked, so this is independent of the large X effect seen). We find no overall difference in Tajima’s D between DE and non-DE loci.

Links between genomic parameters and sex-biased gene expression variation have been a somewhat contentious source of evidence of sexual selection, especially antagonistic forms of sexual selection (Kasimatis *et al*., 2019; Cheng & Kirkpatrick, 2020; Mank *et al*., 2020). Genes that are male-biased in expression show accelerated divergence between species and sex-biased gene expression shows rapid evolution and turnover (Pröschel *et al*., 2006; Harrison *et al*., 2015). Whether sex-biased expression is expected to be related to sex-specific *F*_ST_ or signatures of balancing selection such as Tajima’s D is open to debate, partly because of the potential resolution of antagonistic selection by the strengthening of sex-biased expression. However, there is one very intriguing pattern in our data where the magnitude of change in sex-biased gene expression is related to Tajima’s D. As ΔSB increases (i.e. more male-biased expression in E lines) Tajima’s D in these lines becomes more negative. This pattern is potentially consistent with more resolved sexual conflict in the M lines, because males in M lines are released from sexual selection, and selection driving female-beneficial alleles to high frequency could result in sweeps and/or reduced balancing selection. However, perhaps analyses over the course of the experimental evolution study would be required to convincingly demonstrate associations between changes in sex-bias and potential measures of balancing selection.

In conclusion, we have examined genomic divergence following >160 generations of experimental evolution under altered mating systems. We find that genomic divergence between the experimental lines is highly clustered in the genome, much greater on the X and is associated with changes in gene expression between the experimental lines. Associations with LD and population genetic parameters indicative of selective sweeps or balancing selection are also observed, but are very variable. This raises the possibility that selection has been strong in both M and E lines, but differs in nature (relaxed in M, directional in E), complicating predictions of responses. Overall, our main results support those seen in natural populations, providing an elegant demonstration of the power of experimental evolution to aid the interpretation of complex patterns of natural variation.

## Acknowledgments

This work was part-funded by NERC (grant NE/I014632/1) to MGR & RRS and a combined Natural Environment Research Council and St Andrews 600th Anniversary PhD Studentship grant (NE/L501852/1) to RAWW. Sequencing was supported by the NERC Biomolecular Analysis Facility at the University of Liverpool (NBAF654). Computational analyses were supported by the University of St Andrews Bioinformatics Unit which is funded by a Wellcome Trust ISSF award (grant 105621/Z/14/Z). In addition the authors would like to thank Louise Reynolds, Paulina Giraldo-Perez, and Rudi Verspoor for help with DNA extraction protocols, and Mohamed Noor for helpful advice.

## Author contributions

RAWW performed the data analysis. PV contributed data. The experiment was designed by MGR and RRS. All authors contributed to writing the MS.

## Data Accessibility

Raw reads have been deposited in the short read archive (SRA) of NCBI under the BioProject PRJNA661678

Figures S1 – S7 and Tables S1 – S3 can be found in the Supplementary Material

