## Supplementary Material for "Experimental evolution supports signatures of sexual selection in genomic divergence"

**Wiberg et. al.,**

### Supplementary Material

**Bioinformatic Pipeline**

In all steps outlined below, unless otherwise stated the default parameter values were used. See also the raw reads associated with this study deposited with NCBI (BioProject: PRJNA661678)

**1) Trimming**

Trimming of the reads was carried out using Trimmomatic (v. 0.32; Bolger et al., 2014)

```
$ java -jar trimmomatic
        PE -phred33
        *R1_001.fastq.gz *R2_001.fastq.gz
        *tqc_R1_pe.fq.gz *ftqc_R1_se.fq.gz
        *tqc_R2_pe.fq.gz *ftqc_R2_se.fq.gz
        MINLEN:20 ILLUMINACLIP:${adapters}/TruSeq3-PE.fa:2:30:10
        SLIDINGWINDOW:1:20 MINLEN:20
```

**2) Mapping**

Mapping, indel re-alignment, removal of PCR duplicates, alignment filtering, and merging of .bam files was carried out with bwa mem (v.; Li et al., 2009; Li, 2013) samtools (v. 1.2; Li et al., 2009), GATK (v. 3.3 McKenna et al., 2010; DePristo et al., 2011) and Picard (v. 2.14.1; Broad Institute).

**2.1) Mapping**

```
$ bwa mem -t 5 dpse-all-chromosome-r3.1.fasta *tqc_R1_pe.fq.gz
*tqc_R2_pe.fq.gz > *.sam
$ samtools view -Sb -q 30 -f 0x02 *.sam > *.bam
$ samtools sort -@ 5 -o *_srt.bam *.bam
$ samtools rmdup *_srt.bam *_srt_rmdup.bam
$ samtools sort -@ 5 -o *_srt_rmdup_srt.bam *_srt_rmdup.bam
```

**2.2) Re-alignment around indels**

```
# Add random "readgroup" to .bam
$ java -jar $picard/AddOrReplaceReadGroups.jar
        I= *_srt_rmdup_srt.bam
        O= *_srt_rmdup_srt_rdgrp.bam
        RGID=1
        RGLB=L1
        RGPL=illumina
        RGPU=NONE
        RGSM=[READGROUPNAME]
# Index .bam file
$ samtools index *_srt_rmdup_srt_rdgrp.bam
# Re-align around indels
$ java -Xmx2g -jar $gatk
        -T RealignerTargetCreator
```

```
        -R dpse-all-chromosome-r3.1.fasta
        -I *_srt_rmdup_srt_rdgrp.bam
        -o *_srt_rmdup_srt_rdgrp.intervals
$ java -Xmx4g -jar $gatk
        -T IndelRealigner
        -R dpse-all-chromosome-r3.1.fasta
        -I *_srt_rmdup_srt_rdgrp.bam
        -targetIntervals *_srt_rmdup_srt_rdgrp.intervals
        -o *_srt_rmdup_srt_indraln.bam
# Sort reads and index
$ samtools sort -@ 5 -o *_srt_rmdup_srt_indraln_srt.bam
*_srt_rmdup_srt_indraln.bam
$ samtools index *_srt_rmdup_srt_indraln_srt.bam
# Merge .bam files
$ ls -1 *_srt_rmdup_srt_rdgrp_indraln_srt.bam > in_bam_files
$ bamtools merge -list in_bam_files -out *.bam
$ samtools sort -@ 5 -o *_srt.bam *.bam
```

**2.3) Coverage stats**

```
$ genomeCoverageBed -ibam *.bam -g dpse-all-chromosome-r3.1.fasta> *.cov
```

**3) SNP Calling**

SNP calling was performed with samtools mpileup (v. 1.2; Li et al., 2009) and PoolSNP (Kapun et al., 2018), files were then converted to the .sync format (Kofler et al., 2011).

**3.1) Mpileup**

```
samtools mpileup
        -d 1000000
        -I
        -f dpse-all-chromosome-r3.1.fasta
        R1M1.bam
        R1P1.bam
        R2M2.bam
        R2P1.bam
        R3M1.bam
        R3P2.bam
        R4M1.bam
        R4P1.bam > pseudo_evol.mpileup
```

**3.2) PoolSNP**

```
${PoolSNP}/PoolSNP.sh mpileup=pseudo_evol.mpileup
        output=pseudo_evol
        reference=dpse-all-chromosome-r3.1.fasta
        names=R1M,R1P,R2M,R2P,R3M,R3P,R4M,R4P
        min-cov=17
        max-cov=0.95
        min-count=16
        min-freq=0.001
        miss-frac=0.01
        base-quality=15
        jobs=10
```

**4) QBGLM Analysis**
QBGLM analysis was performed with scripts from Wiberg *et al.,* (2017), available from github (https://github.com/RAWWiberg/poolFreqDiff)

**Permutation tests**

To further test the QBGLM method under a "null" scenario, we performed a permutation test to determine the expected number of "top SNPs" (SNPs with q-values < 0.05) as well as the level of clustering of such "null" SNPs. We permuted the E and M labels among all 8 samples and consider 10 different permuted datasets. With all treatment labels randomly assigned to the SNP data, we never observe as many "top SNPs" as we found in the real data. In the permuted datasets we observe between 2 and 107 (out of 1,852,324) SNPs with q-values < 0.05. These SNPs give a correspondingly smaller number (between 1 & 32) of "clusters" (defined in the same way as in the manuscript). Thus we have substantially more outliers in the observed dataset than in the permuted ones. Clustering of SNPs arises, in part, due to the closely linked SNPs having correlated allele frequencies and therefore showing similar results. This will be true whether change is due to genetic drift or due to selection, especially, relative to evolutionary time, over only a small number of generations. Thus taking the 480 most differentiated SNPs from the permuted datasets above unsurprisingly produces a similar number of clusters as in the observed data (between 8 and 77), because the permutations are done in such a way as to preserve this correlation structure of allele frequencies at closely linked SNPs. However, most of these 480 "pseudo-top" SNPs have q-values of ~0.5 indicating that they do not show consistently different allele frequencies between E and M lines.

We further explored the appearance of "clustering" in a smaller test dataset using 1,000 random SNPs where we conducted a second permutation test which removed this correlation structure among closely linked SNPs by permuting E and M labels for each SNP independently rather than for the SNP set as a whole. We then compared results for the same set of 1,000 SNPs but where the labels are permuted for the entire set (preserving the linkage effect). We performed this procedure 10 times both ways. When removing the correlation we observe between 11 and 24 SNPs with p-values < 0.05 (mean = 16.1, out of 1000 SNPs). If we maintain the correlation structure we obtain between 9 and 18 SNPs with p-values < 0.05 (mean = 15.4). However, as expected, if we use the top 100 SNPs for clustering, then we see more "clusters" (between 87 and 93 mean = 90.8) when the correlation structure is removed than when it is maintained (87-95, mean = 86.7). That is, when allele frequencies are not allowed to be correlated, the significant SNPs are more evenly spread throughout the genome. Therefore the correlation between allele frequencies in the observed data is contributing to the "clustering" of SNPs.

**5) Functional Analysis**
Functional analysis was carried out with GOwinda (Kofler & Schlötterer 2012), the AME tool from the MEME package (Bailey et al., 2009; McLeay & Bailey 2010). Closest genes to top SNPs were identified with bedtools (v. 2.26 Quinlan & Hall, 2010).

**4.1) Gowinda**

```
$ java -Xmx4G -jar ~/bin/Gowinda-1.12.jar
        --snp-file *all_snps.tab
        --candidate-snp-file *fixed_snps.tab
        --annotation-file Dpse_genes_dmelnames.gtf
        --gene-set-file dmel_funcassociate_go_associations_mod.txt
        --output-file *GOwinda.out
        --simulations 1000000
        --gene-definition updownstream1000000
```

**4.2) MEME**

```
$ ame --oc ame_out --pvalue-report-threshold 1 --control
*all_SNPs_regions.fasta *SNPs_regions.fasta motifDataBase
```

### Supplementary Tables and Figures

**Table S1.** List of R packages and references.

| Package | Version | Citation/Source |
|---|---|---|
| plyr | 1.8.4 | Wickham H (2011). The Split-Apply-Combine strategy for data analysis. *Journal of Statistical Software.* **40:** 1-29. |
| dplyr | 0.50. | Wickham H (2011). The Split-Apply-Combine strategy for data analysis. *Journal of Statistical Software.* **40:** 1-29. |
| scales | 0.4.1 | Wickham H (2016). scales: scale functions for visualization. https://CRAN.R-project.org/package=scales |
| reshape2 | 1.4.2 | Wickham H (2007). Reshaping data with the reshape package. *Journal of Statistical Software.* **21:** 1-20. |
| stringr | 1.2.0 | Wickham (2017). stringr: Simple, consistent wrappers for common string operations. https://CRAN.R-project.org/package=stringr |

**Table S2.** Coverage and mapping statistics. Given are the total reads that were mapped, the proportion of reads that mapped and passed all filters, and the mean and median coverage.

| Sample | Total Reads | Mapped Reads (%) | Coverage Median [Mean] |
|---|---|---|---|
| R1M | 33,750,918 | 27,782,976 (82) | 33x [32.2x] |
| R1E | 34,284,906 | 28,619,221 (84) | 35x [33.2] |
| R2M | 32,142,578 | 26,622,688 (83) | 32x [30.8] |
| R2E | 38,020,312 | 31,319,496 (83) | 38x [36.3] |
| R3M | 27,025,159 | 22,213,842 (82) | 26x [25.9] |
| R3E | 37,764,199 | 30,823,152 (82) | 37x [35.7] |
| R4M | 29,802,499 | 24,901,734 (84) | 30x [28.9] |
| R4E | 31,449,467 | 25,068.166 (80) | 30x [29.1] |

**Table S3.** Distribution of chromosome lengths, the overall number of SNPs and the number of outlier SNPs across the main chromosome arms.

| Chromosome | Length | N SNPs[a] | N top SNPs[b] |
|---|---|---|---|
| 2 | 30,819,483 | 511,005 | 30 |
| 3 | 19,787,792 | 169,950 | 200 |
| 4 | 27,243,186 | 489,294 | 30 |
| XL | 24,770,255 | 320,645 | 112 |
| XR | 24,741,483 | 264,625 | 108 |
| Total on Chromosomes | 127,362,199 | 1,755,519 | 480 |
| Overall Total | - | 1,852,324 | |

*a.* There are excesses (compared to those expected from the relative lengths of the chromosomes) on chromosomes 2 and 4 (Chi-squared = 109,095, d.f. = 4, $p < 0.001$).

*b.* There are excesses (compared to those expected from the relative lengths of the chromosomes) on chromosomes 3 and the X chromosome arms (Chi-squared = 698.26, d.f. = 4, $p < 0.001$).

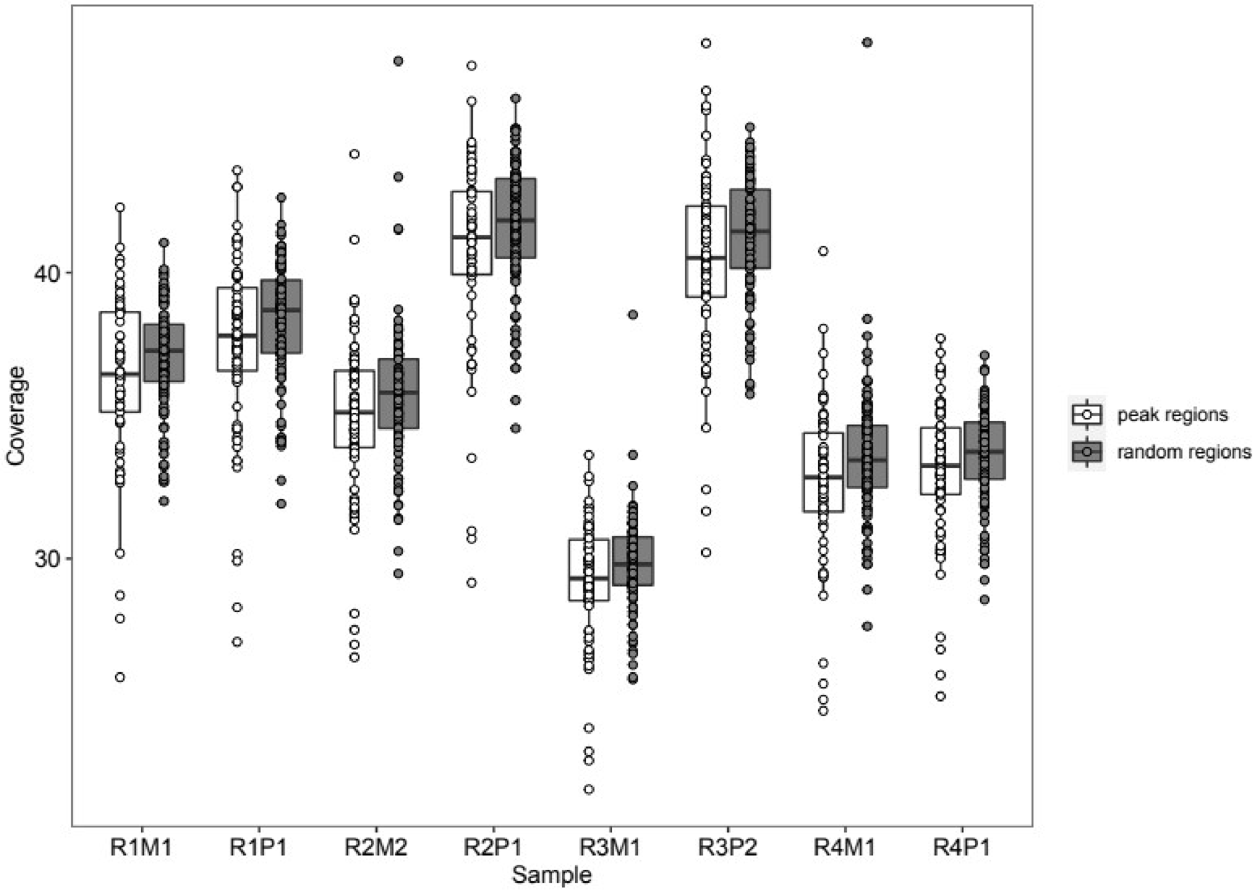


**Figure S1.** Coverage distributions between the regions around top SNPs (peak regions) and 100 randomly sampled regions with a similar length distribution (random regions). In an ANOVA, the difference between peak regions and random regions is significant ($F_{(1,1351)} = 34.5$, $p < 0.001$), but the effect of sample is much larger ($F_{(7,1351)} = 459.1$, $p < 0.001$).

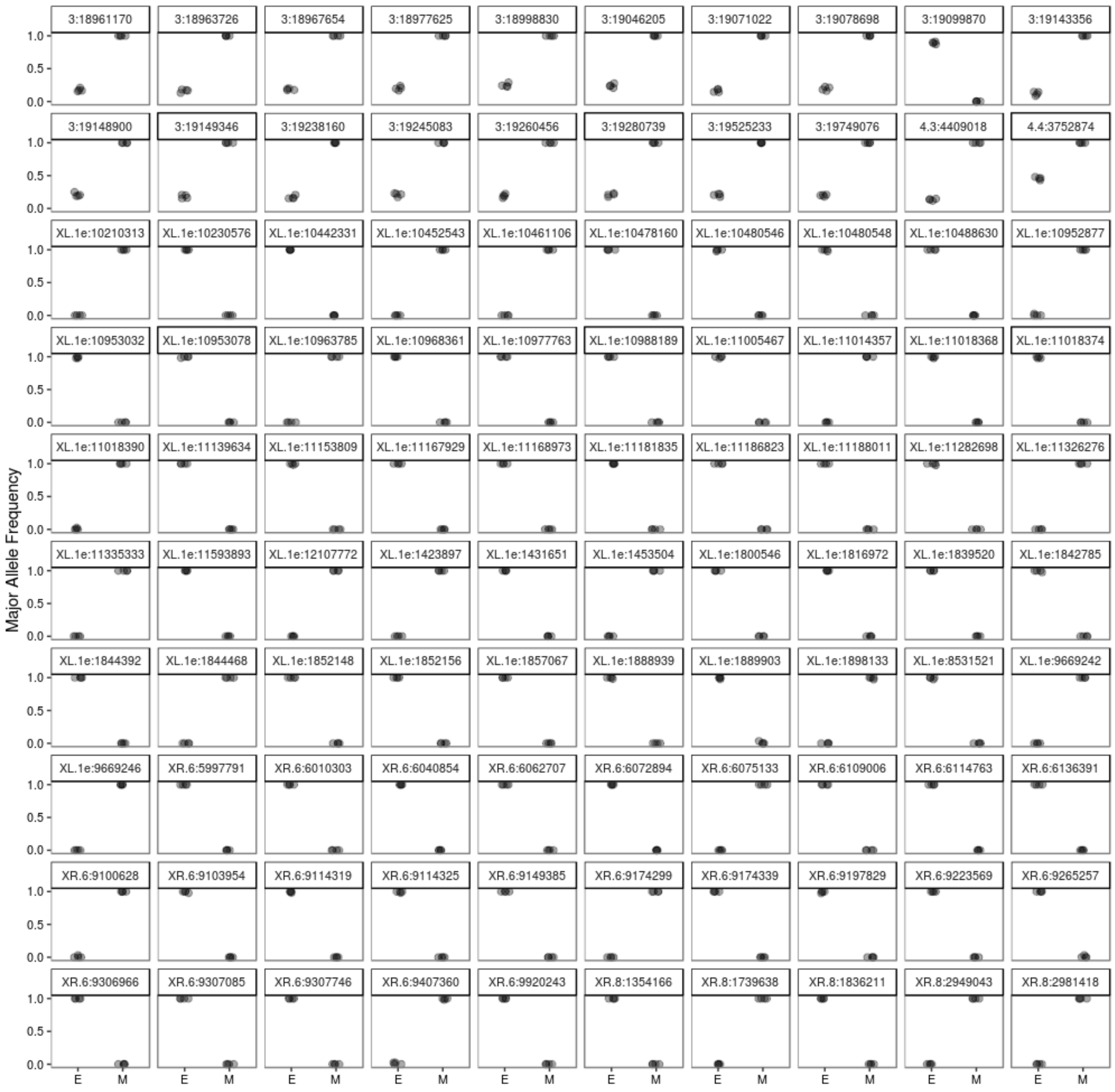


**Figure S2.** The allele frequencies in E and M lines for the top 100 SNPs with the lowest q-values from a quasibinomial GLM of allele frequency differences. The numbers in the panel titles give the chromosome and the position along the chromosome of each SNP.

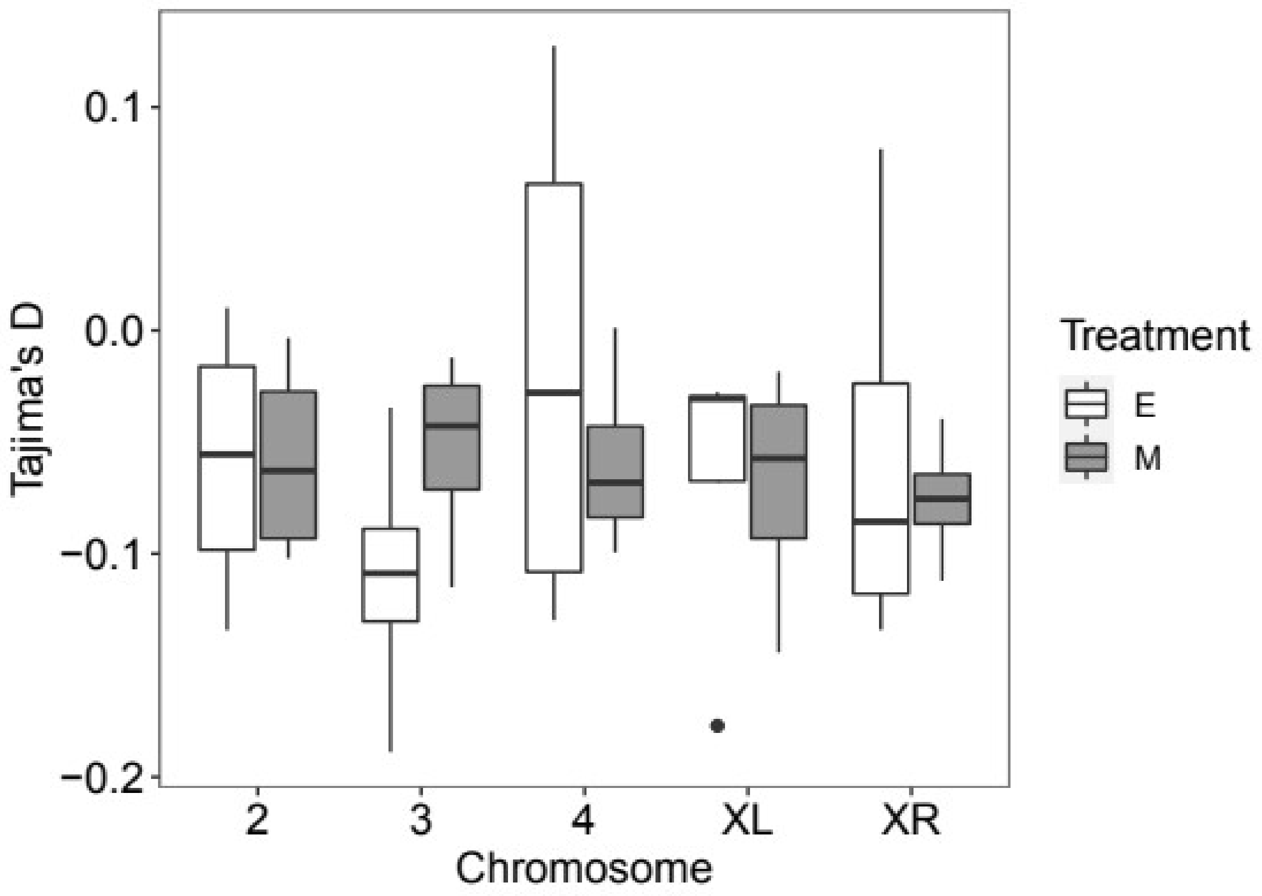


**Figure S3.** Levels of genetic diversity (Tajima's D) on each chromosome in E and M lines. Tajima's D) is estimated in overlapping windows of 50kb, then averaged across the chromosomes. Boxplots show the distribution of Tajima's D on each chromosome across replicate experimental evolution lines.

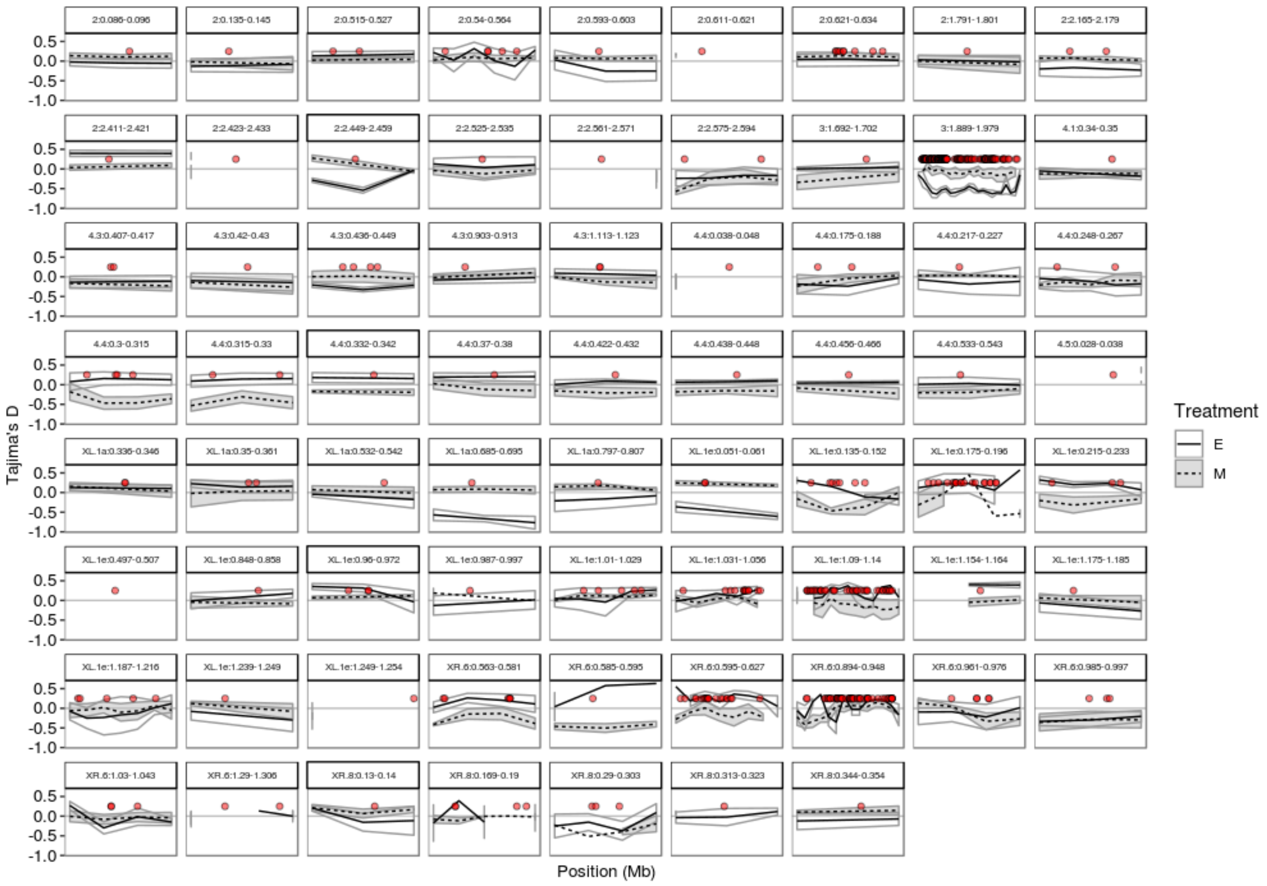


**Figure S4.** Mean (±SE) of Tajima's D in overlapping 50 kb windows along the chromosomal reigons underneath the 70 peaks of highly differentiated SNPs. Panel titles give the chromosome and start and end positions (in Mb) of the regions. X-axis tickmarks have been removed for clarity but note that the scale changes across panels. Red points denote the locations of SNPs with q-values < 0.05, these have been plotted at the same coordinates on the y-axis for convenience.

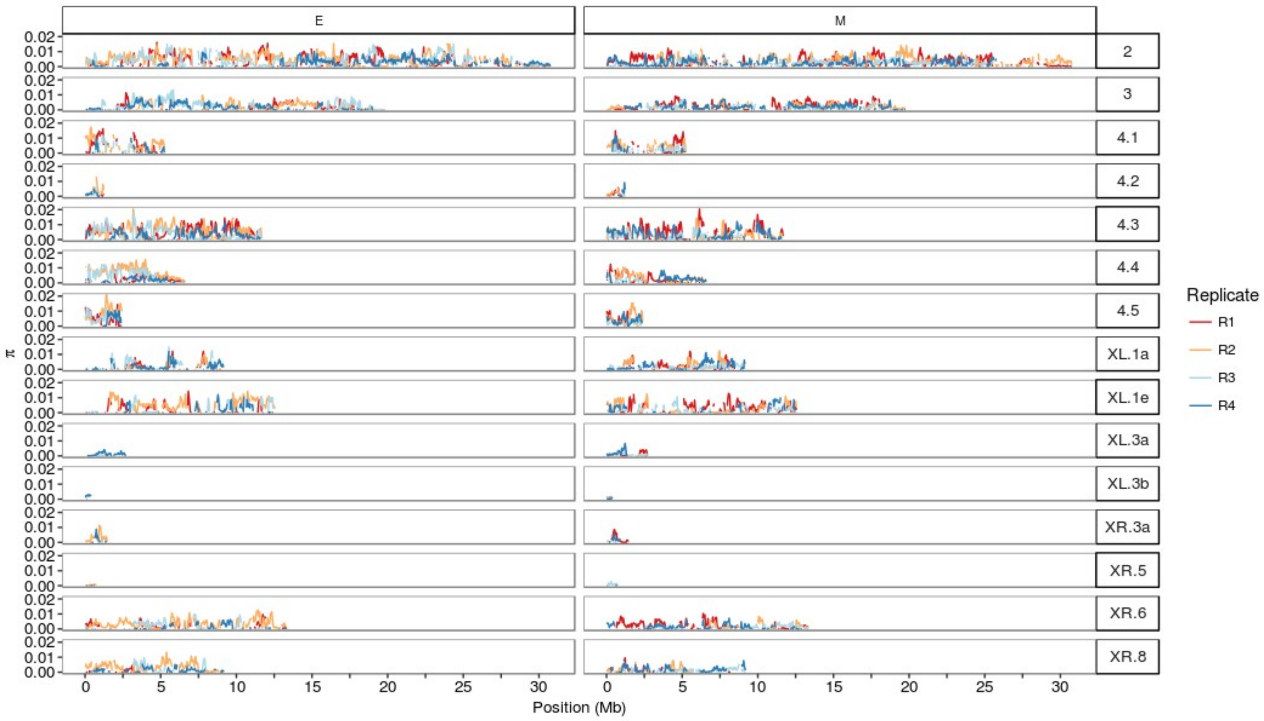


**Figure S5.** π in overlapping 50kb windows across chromosomes and replicates of E and M lines.

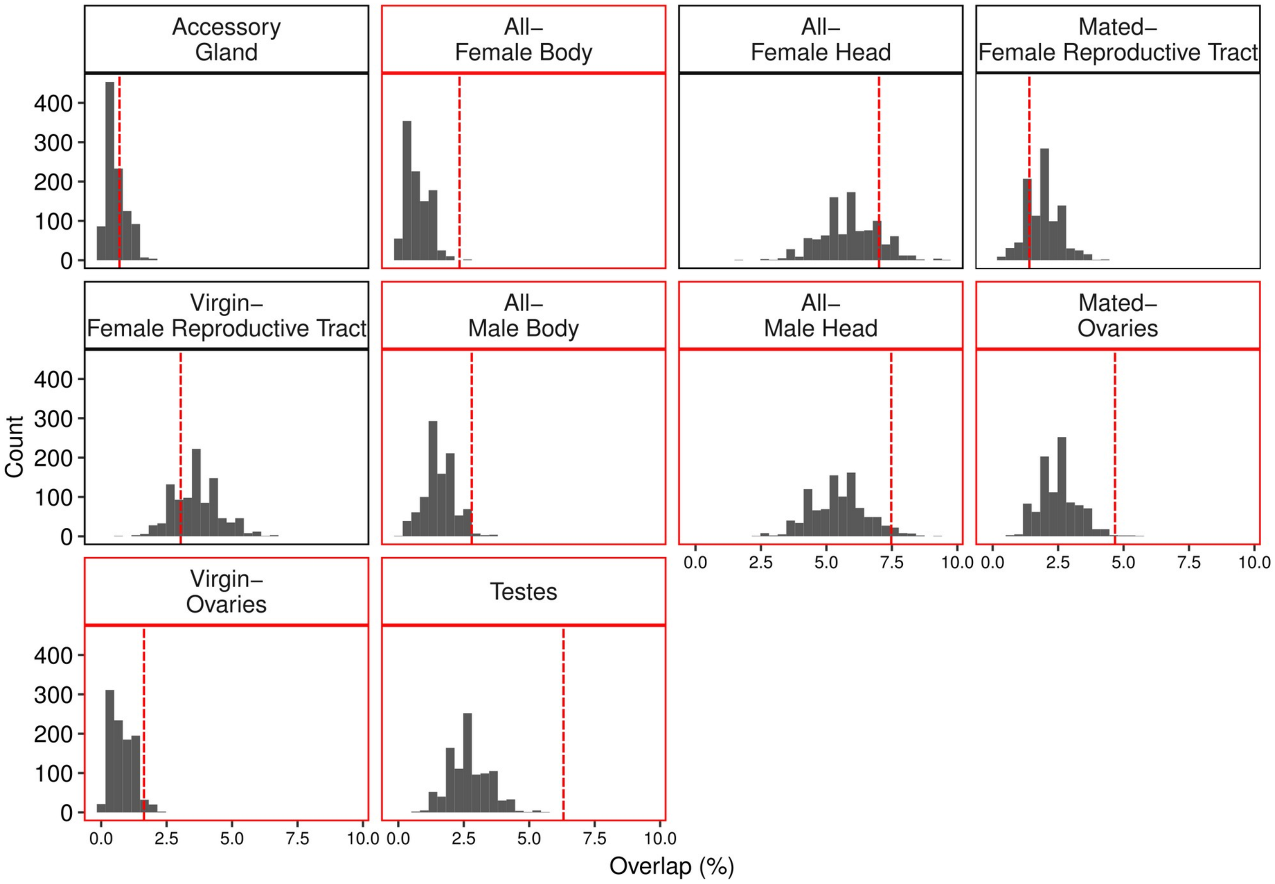


**Figure S6.** Distributions of overlap between bootstrap samples of genes and sets of differentially expressed genes from Veltsos et al., (*unpublished*). Vertical dashed lines indicated the empirical overlap of the 428 genes within 10kb of top SNPs for each set. Panels highlighted in red have an empirical overlap >= the 95$^{th}$ percentile of the bootstrap distribution.

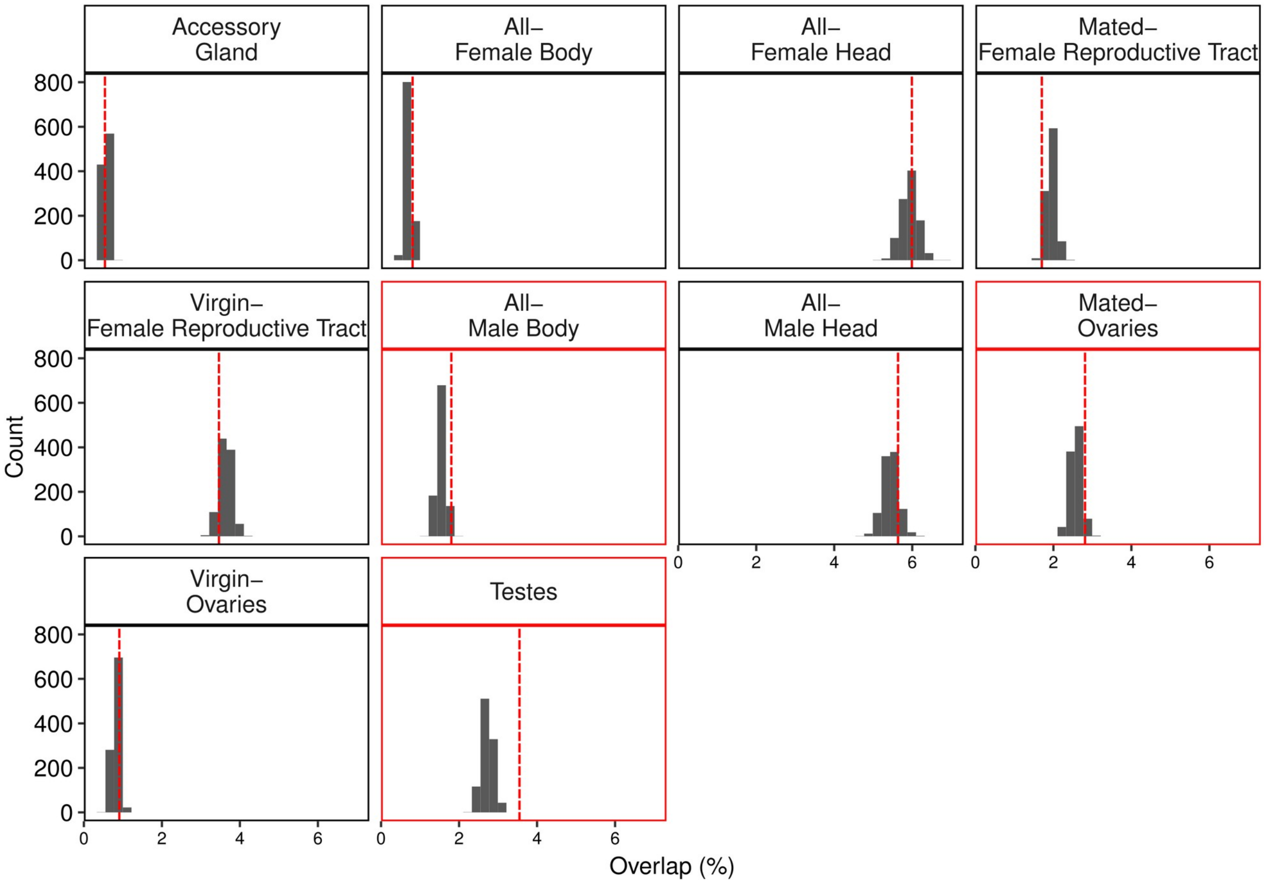


**Figure S7.** Distributions of overlap between bootstrap samples of genes and sets of differentially expressed genes from Veltsos et al., (*unpublished*). Vertical dashed lines indicated the empirical overlap of the 7,045 genes within 1Mb of top SNPs for each set. Panels highlighted in red have an empirical overlap >= the 95th percentile of the bootstrap distribution.
